# Neural representations of vicarious rewards are linked to interoception and prosocial behaviour

**DOI:** 10.1101/2022.03.04.482889

**Authors:** Luis Sebastian Contreras-Huerta, Michel-Pierre Coll, Geoffrey Bird, Hongbo Yu, Annayah Prosser, Patricia L. Lockwood, Jennifer Murphy, Molly J. Crockett, Matthew A.J. Apps

## Abstract

Every day we constantly observe other people receiving rewards. Theoretical accounts posit that vicarious reward processing might be linked to people’s sensitivity to internal body states (interoception) and facilitates a tendency to act prosocially. However, the neural processes underlying the links between vicarious reward processing, interoception and prosocial behaviour are poorly understood. Previous research has linked vicarious reward processing to the anterior cingulate gyrus (ACCg) and the anterior insula (AI). Can we predict someone’s propensity to be prosocial or to be aware of interoceptive signals from variability in how the ACCg and AI process rewards? Here, participants monitored rewards being delivered to themselves or a stranger during functional magnetic resonance imaging. Later, they performed a task measuring their willingness to exert effort to obtain rewards for others, and a task measuring their propensity to be aware and use interoceptive signals. Using multivariate similarity analysis, we show that people’s willingness to be prosocial is predicted by greater similarity between self and other representations in the ACCg. Moreover, greater dissimilarity in self-other representations in the AI is linked to interoceptive propensity. These findings highlight that vicarious reward is linked to bodily signals in AI, and foster prosocial tendencies through the ACCg.

## Introduction

From seeing strangers enjoy delicious-looking meals in restaurants, to observing likes on social media posts, witnessing other people receive rewards is a fundamental feature of our social lives. Theories suggest that the vicarious processing of these rewards is a key component of social behaviour (1, 2). More strongly representing others’ rewards may allow people to share others’ positive experiences and lead to stronger visceral responses (2–7), as well as it may lead us to choose to exert effort into prosocial acts aimed at obtaining positive outcomes for them (8–12). As such, representing other people’s outcomes may be linked to the propensity to be aware of internal bodily signals that influences our social behaviour. However, largely, such accounts have not been empirically tested. Although some previous studies have linked vicarious reward processing to prosocial behaviour (13–15), and variability in interoception to the processing of social information (5–7, 16, 17), it is unclear whether it is the vicarious representation of others’ rewards that is linked to interoception and prosocial behaviour, or some other aspect of the tasks that are used to probe them (e.g. the desire to help). As a result, the neural mechanisms that relate either interoception or prosocial behaviour to vicarious reward signals remain poorly understood.

Research examining self and vicarious reward processing has implicated two regions of the brain as consistently responding to self and other rewards - the gyral portion of the anterior cingulate cortex (ACCg) and the anterior insula (AI) (2, 18–21). These specific portions of the cingulate and the insular cortices have been also largely associated with vicarious processing and social stimuli in general, suggesting a specialised function in socially-relevant stimuli, which contrasts with other portions of these regions (e.g. ACC sulcus, posterior insula) more focused on processing self events (2, 18, 29–31, 21–28). Importantly, among these functions, these two regions are engaged when we see cues indicating other people will receive a reward, respond differently when people receive a reward themselves, and do not respond to foregone rewards delivered neither to self or other (2, 21). Strikingly, separate lines of research also implicate these same regions in interoceptive processes and to prosocial behaviour (15, 25, 32–34). As a result, theoretical accounts suggests that reward processing, interoception and motivation may be intimately linked (35–37). However, it is unclear whether vicarious reward processing in the ACCg and AI are linked separately to these different functions.

Indeed, research has suggested that despite both the ACCg and AI being engaged when seeing others’ rewards, the processing in each region may serve different functional roles. Broadly speaking, the insula is engaged in processing the links between information about rewards and bodily states, while ACC may be more strongly linked to driving the motivation to obtain rewarding outcomes (36–45). AI has been proposed as key in processing interoceptive signal, self-awareness and body ownership, which might be crucial to disentangle vicarious from self outcomes (34, 38, 46). On the other hand, ACCg has been involved in prosocial motivation and vicarious processes, with some studies reporting specific responses for social stimuli, while others showing general activity for self and other (2, 18, 21–26, 47, 48). However, as most tasks used to examine the neural processes underlying prosocial behaviour involve self and other rewards, and also may evoke interoceptive processes, it has been difficult to disentangle whether vicarious processing in the ACCg and AI is associated with distinct functions, and how specific or general these roles are for self and vicarious processes. We propose one solution to this problem is to examine whether neural responses to others’ rewards in the ACCg and AI when simply passively observing them are linked to either individual differences in people’s willingness to be prosocial, or their propensity to be aware of interoceptive signals, in separate tasks.

How can we interpret neural responses to others’ rewards? While a number of studies have examined vicarious reward processing with functional magnetic resonance imaging (fMRI), there are contrasting viewpoints on how vicarious reward BOLD signals should be interpreted (1, 2, 25, 47). On the one hand, the overlap between clusters that respond to self and other rewards in the ACC and AI is often interpreted as being indicative of a ‘common currency’ that represents information about both ourselves and other people in the same manner (1, 49). Thus, it is assumed that this neural overlap could be indicative of a similar positive affective response at another receiving rewards to rewards being received by oneself. In contrast, other studies have shown that greater specialisation – more distinct processing between self and other rewards – is linked to higher levels of empathy and higher learning speed for prosocial actions (20, 50). Such greater specialisation, rather than a common currency, might therefore underlie stronger prosocial tendencies (20, 47, 50). However, only a few studies have examined variability in vicarious reward processing between people, with the majority of them using univariate fMRI analysis, where the strength of activity in a certain brain location is correlated between self and other conditions. Even though these studies have provided valuable evidence to support either perspective on this dispute, there is an inherent limitation of this approach - similar strength in univariate BOLD signal does not necessarily imply similar multivariate patterns of responses for self and other rewards. Therefore, even in the presence of neural overlap between self and other rewards in some individuals, there could be a different pattern of response for each condition within a brain area, suggesting distinctive processing. Crucially, analysis techniques that can formally test the similarity between self and other reward representations, such as multivariate pattern analysis, have rarely been employed in previous studies, and typically without paying attention to individual differences.

Here, we use a multivariate approach to test whether similarity in neural responses to self and other reward in the AI and ACCg is linked to people’s propensity to be aware and use interoceptive signals and their willingness to exert effort into prosocial behaviours. Participants performed three tasks (**Fig. 1**), the first while undergoing fMRI and the second two in a separate behaviour session. Participants (n = 61) first underwent fMRI during a value representation task, in which they observed cues indicating that rewards were being accrued – or being taken away – from oneself or from an anonymous other person. In addition, they performed two behavioural tasks without fMRI. First, an interoceptive respiratory task, which measured people’s propensity to be aware and use internal, rather than external, signals to judge their respiratory output (51). Second, a prosocial effort task (8), which measured willingness to exert physical effort to obtain rewards for oneself or for an anonymous stranger (8, 10, 48, 52). These two tasks enable measurement of individual differences in reliance on interoceptive cues and prosocial motivation. To examine whether the distinctiveness of self and other reward representations in ACCg and AI is linked to prosocial motivation and interoception respectively, offline task behaviour was correlated with pattern similarity in each region from the value representation task (53).

**Figure 1.**
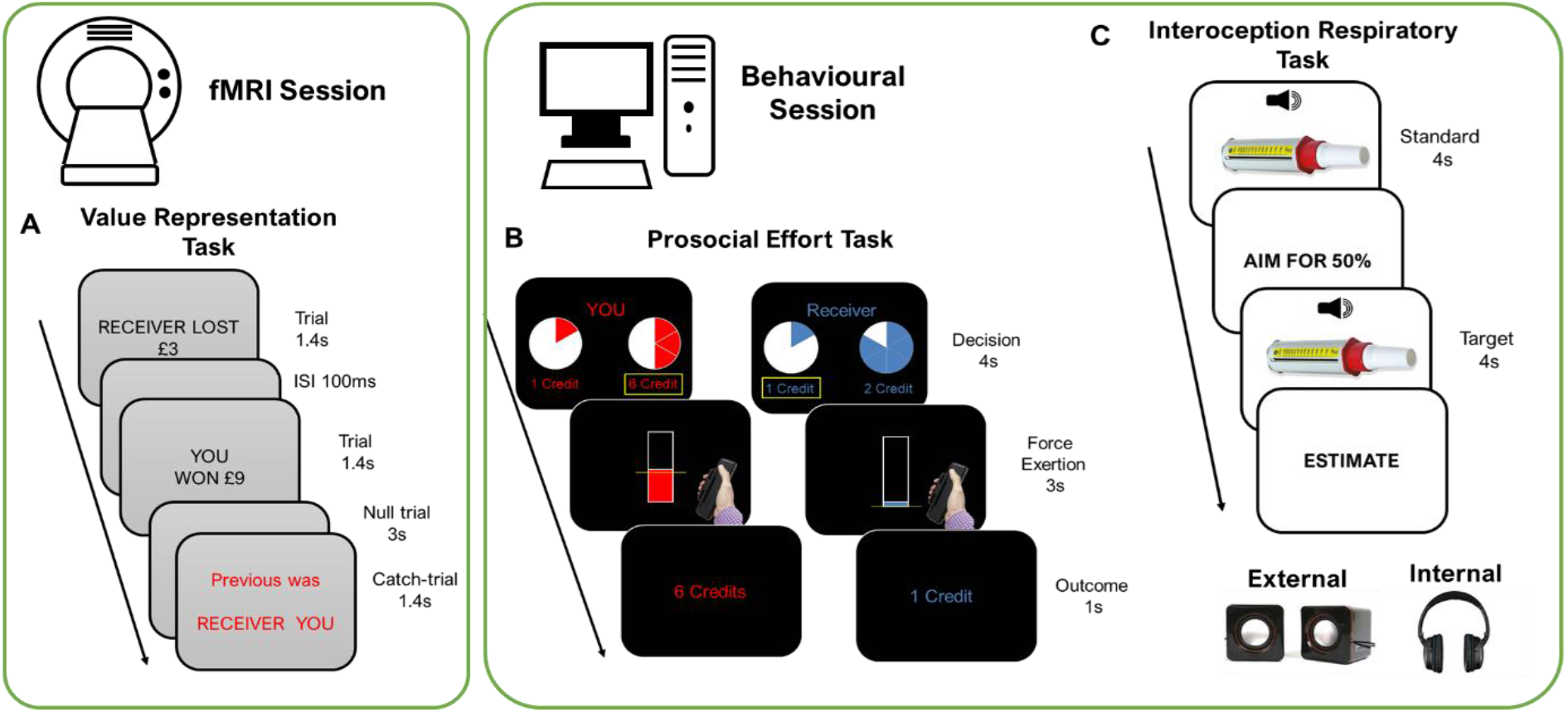
fMRI and Behavioural tasks. Participants completed three tasks across two sessions separated by at least one week A. *fMRI session: The value representation task*. This task measured neural similarity between representation of rewards and losses for the self and others. Participants passively witnessed financial gains (6 levels) and losses (6 levels) for themselves and for another unknown person, the Receiver. Catch trials asked either who was the recipient of the last outcome, or the magnitude of the reward on the previous trial, to encourage attention to the task. Similarity was calculated as the Pearson correlation between the spatial representation of self- and other-related neural activity. *B Prosocial Effort task*. Participants made choices about whether to exert different amounts of effort (30-70% of their own maximum grip strength) on a handheld dynamometer for variable amounts of reward (2-10 credits). Participants worked either to benefit themselves or an anonymous other. *C. Interoceptive Respiratory Task*. On each trial participants blew into a peak-flow meter twice, the first blow setting a standard for that trial. In the second blow, participants were required to achieve a percentage of the first, with participants estimating their actual percentage performance at the end of the trial. This was conducted in an internal condition with white noise played through headphones to prevent the use of external auditory cues, and an external condition where external auditory cues were available. The difference in estimation accuracy between internal and external conditions was taken as a measure of how much participants rely on internal vs external signals. Note that the sequence of screens is an illustration of the test, as participants were blindfolded.

We find that prosocial motivation and reliance on interoceptive signals are linked. The more strongly guided by internal signals someone is, the more incentivised they are to obtain rewards for another person. Strikingly, people’s levels of prosocial motivation and reliance on internal signals are related to multivariate neural patterns in the ACCg and AI, respectively. More similar patterns between self and vicarious reward in the ACCg are associated with increased levels of prosocial motivation but not motivation to obtain rewards for oneself. However, more dissimilar patterns in the AI are correlated with higher propensity to be aware and use interoceptive signals. These results highlight that even when simply observing other’s rewarding outcomes, variability in responses in the ACCg and AI can be linked to subsequent levels of prosociality and interoception, respectively.

## Results

During the first fMRI session, participants completed a value representation task, where on separate trials either themselves, or an anonymous stranger received or lost rewards of different magnitudes. The stranger was a confederate who participants believed to be another participant in the experiment. We followed a rigorous protocol (8, 54) to make sure participants believed in the existence of the other person, ensuring anonymity and confidentiality (see **Materials and Methods** and **Supplementary Information** for details).

In the behavioural session, the prosocial effort task required participants to make choices of whether to squeeze a grip force device to obtain rewards either for themselves or for an anonymous other. This task measured the amount of reward necessary to incentivise people to work, and the degree of avoidance of the required physical effort when deciding whether to work for oneself or for another person. In this behavioural session, participants also completed the respiratory interoception task, where in each trial they performed an exhalation into a standard peak flow meter, and then a subsequent second exhalation aiming to achieve a target percentage of the first exhalation. After the second exhalation, participants were required to report what percentage they actually achieved. The difference between their estimation and the actual percentage of their exhalation served as an index of accuracy (see **Materials and Methods equation 1**). Trials were performed in one of two conditions, an ‘internal’ condition where no external auditory cue could guide their performance, and an ‘external’ condition where external auditory cues caused by their exhalation were available to be used. Using this technique, a metric of reliance on interoceptive cues was calculated, by comparing estimation accuracy between the internal and external conditions (see **Materials and Methods equation 2**). If participants rely on exteroceptive cues, then removal of those cues should hamper performance; conversely, if participants rely on interoceptive cues then performance should be unaffected by removal of external cues. On this task, therefore, more positive values reflect more of a reliance on (i.e. more propensity to be aware and use) interoceptive cues to complete the task (**Supplementary Fig. S1A**).

### Behavioural results: reliance on interoceptive signals is associated with prosocial motivation

Previous research has suggested that people’s propensity to rely on interoceptive signals is correlated with their levels of altruism (16, 55). However the tasks used to measure prosocial or interoceptive tendencies may confound different processes together and there have been questions of their validity (51, 56, 57). Here, we address these issues using tasks to measure prosocial behaviour and interoceptive processes without such confounds. Firstly, studies have used economic games in which the rewards delivered to the other person directly impact on the magnitude of rewards obtained oneself. Although interesting for quantifying variability in one’s desire to benefit others, they cannot distinguish between two different motives - sensitivity to one’s own rewards, or an increasing desire to give rewards to other people (11, 47). Here, we use a prosocial effort task that is able to measure sensitivity to one’s own and other’s rewards, as well as separately measures sensitivity to exerting effort for oneself and others. Secondly, we used a respiratory interoceptive task that measures individual differences in how much people rely on interoceptive information, i.e. the propensity of people to be aware and use bodily signals, that has been validated and have overcome some of the problems raised on previous tasks (51, 58). Furthermore, this task focuses on the extant in which people are aware of their internal signal, rather than simply interoceptive accuracy, which could be directly related to the role of AI in self-awareness and body ownership (34, 38). Thus, using these two tasks we can therefore examine the relationship between interoception and prosocial tendencies and test the hypothesis that stronger sensitivity to other’s rewards is linked to one’s reliance on interoceptive processes

We tested whether reliance on interoceptive signals was associated with motivation for self or/and others’ rewards. A mixed effects model predicting choices to work or rest in the prosocial effort task was conducted including level of effort, magnitude of reward, and beneficiary (self vs other), together with their interactions (up to three-way), as fixed effects predictors. Random effects were also included, with random intercepts at the subject level, and random slopes on effort level and reward magnitude (see **Materials and Methods equation 3**). Model comparison revealed that including participants’ interoception scores from the respiratory task as an additional between-subjects predictor of choices in the prosocial effort task (see **Materials and Methods equation 4**) improved model fit, both when penalising the model for additional parameters (AIC: simple model = 5115.5, interoception model = 5079.3; **Supplementary Fig. S1B**) and when performing a ratio test on model log-likelihoods (χ^2^_diff_ = 52.28, df_diff_ = 8, p < 0.001). Thus, decisions to work in the prosocial effort task were better explained by a statistical model that included participants’ reliance on internal signals.

We hypothesised that people who relied more on internal signals would be more motivated to act prosocially, and that this would be specifically linked to how incentivised people are by others’ rewards. Consistent with this, we found a three-way interaction between interoception score, reward and beneficiary in the mixed effects model (b = 0.42, SEM = 0.11, z = 3.87, p < 0.001, **Fig. 2A** and **2B**). Thus, during self trials participants were more likely to work as the rewards on offer increased, but this effect was present regardless of their interoception score (**Fig. 2A**). In the other (prosocial) trials, there was an effect of interoception, whereby people who relied more on interoceptive signals were more incentivised to work at higher reward levels for others. Conversely, externally-focused people were not incentivised to choose to work more often as the rewards that would be received by the other person increased (**Fig. 2B**).

**Figure 2.**
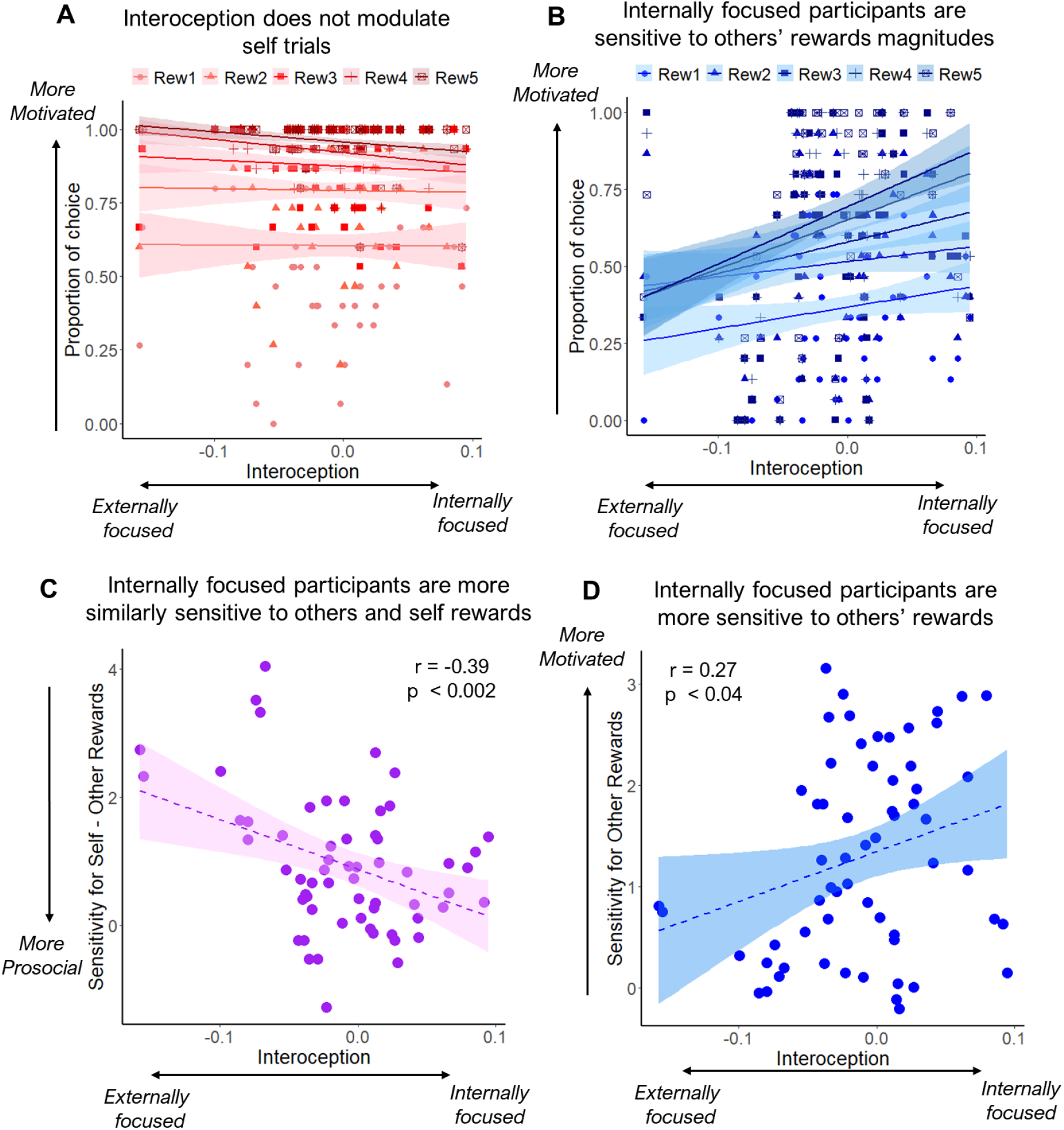
Reliance on interoceptive signals is associated with prosocial choices. A. Interoception score did not influence motivation for rewards in self trials. Participants did not show different patterns in their proportion of choices to work over rest (y-axis) depending on their level of reliance on internal vs external signals (x-axis) across different reward magnitudes. B. Interoception influences motivation for others’ rewards. As participants relied more on internal than external cues, they chose to work more as the reward to be received by another person increased. Participants who relied more on external cues were more reluctant to work to benefit others regardless the reward on offer. Shaded areas show the 70% confidence interval around the slopes. Individual points show the score of each participant for each condition. C. People who rely more on internal vs external signals (x-axis), weighted rewards more similarly for self and other when making decisions to work to obtain rewards. Y-axis depicts the difference between self and other betas from two mixed models predicting choices separately for self and other. Higher values indicate self rewards are valued more than others’. D. Participants who relied more on internal vs external cues (x-axis), were more sensitive to others’ rewards (beta estimates, y-axis). Shaded areas show the 95% confidence interval around the slopes. Individual points show the score of each participant.

In addition, there was a two-way interaction between interoception and beneficiary (b = 0.65, SEM = 0.12, z = 5.3, p < 0.001), such that participants who relied more on internal than external signals were more willing to work for others regardless of the rewards on offer or effort required (**Supplementary Fig. S2A**). Notably, despite previous research suggesting a link between interoception and people’s sensitivity to effort, there was no such interaction in our data (see **Supplementary Table S1** for a full description of these results). Finally, the remaining findings of the mixed model replicated the results of previous studies using the prosocial effort task (8, 10, 48, 52), with significant effects of effort, reward and beneficiary, as well as interactions between reward and beneficiary, and effort and beneficiary (see **Supplementary Fig. S3A** and **S3B**). In sum, we show that how much someone relies on internal signals is linked to how prosocial they are and not to their motivation to work for themselves. Specifically, more internally-focused individuals are more incentivised by the rewards that can be obtained for another person, and work more when they can obtain a bigger benefit for them.

Given that in the prosocial effort task the majority of people choose to work more at lower reward levels for themselves than for other people, this leads to a prediction that people with more positive interoception scores (i.e. more internally focused, relying more on interoceptive signals) would show less of a difference in decisions to work between self and other, as a function of reward. Furthermore, given the results of the mixed model above, greater interoception scores should be specifically associated with sensitivity to others’ rewards and not with sensitivity to self rewards nor sensitivity to effort. Thus, as a confirmatory, post-hoc analysis, we extracted beta parameters for effort and reward from two mixed effect models where decisions to work or rest were taken separately for self and other trials (see **Materials and Methods equations 5 and 6**). Thus, these indexes were a proxy for individual differences in sensitivities to effort levels and magnitudes of rewards for self-benefitting and prosocial decisions.

From the results described above, we expected interoception to be associated with sensitivity to others’ rewards and not to effort or self rewards. We tested this using Pearson correlations between the interoception score, and reward and effort betas for self, other and their difference. Results revealed a significant negative correlation between interoception and the difference in reward betas, such that people who relied more on internal signals were more similarly motivated for self and other rewards, while people who were more externally focused valued more their own rewards compared to others’ (r_(59)_ = −0.39, p < 0.003, **Fig. 2C**). To test if this effect of interoception on prosocial motivation was specific to sensitivity to rewards, we correlated difference in effort betas with interoception. We found null correlation (r_(59)_ = 0.15, p = 0.26), which was significantly different from the correlation between interoception scores and sensitivity to reward (Fisher’s Z transformation, z = - 3, p < 0.003). As expected, the interoception score also correlated with sensitivity to others’ rewards (r_(59)_ = 0.27, p < 0.04, **Fig. 2D**), but not with self rewards (r_(59)_ = −0.14, p = 0.29; **Supplementary Fig. S2B**), and these effects were significantly different from each other (z = 2.23, p < 0.03), suggesting specificity of the social effects. Furthermore, interoception did not correlate with neither other (r_(59)_ = - 0.08, p = 0.54) nor self effort betas (r_(59)_ = −0.13, p = 0.32), and these correlations showed a significant difference (z = 2.19, p < 0.03) and a non-significant trend from its effects on other reward betas (z = 1.92, p = 0.055), suggesting that interoception is specifically associated to sensitivity to others’ rewards in the effort task. Taken together, these results reveal that people who rely more in internal versus external signals are specifically more driven by others’ rewards when deciding whether to expend energy to act prosocially.

### fMRI results: Different roles of the ACCg and the right dorsal AI in vicarious rewards

Next, we examined whether the degree of similarity between neural patterns evoked by self and other rewards during the fMRI value representation task was predicted by prosocial motivation and interoception. Multivariate similarity analysis was calculated as the Pearson correlation between the average parametric map for self and other rewards (53). Thus, the similarity index represents the spatial similarity between the patterns of activation for self and other rewards across the different reward levels for each participant. Our hypotheses specifically related to the ACCg and AI due to their unique profiles of being linked to interoception, prosociality and vicarious reward processing (2, 15, 19, 25, 32, 38). We therefore included five ROIs: four AI portions - right dorsal AI (RdAI), right ventral AI (RvAI), left ventral AI (LvAI) and left dorsal (LdAI) -, and the ACCg (bilateral). Traditional whole-brain univariate analyses performed on the same data are reported in **Supplementary Tables S2 and S3**.

Five regression models were used to test whether the degree of similarity between self and other representations of reward was associated with interoception, one in each ROI, with the similarity of self and other representations being the dependent variable and participants’ interoception score (reflecting their reliance on interoceptive signals) as a predictor. Only one area, the RdAI, showed a significant effect (b = −1.01, F_(1,52)_ = 11.12, p < 0.008 FDR corrected). No effects were found in any of the other ROIs. Within the RdAI, less similar responses between self and other rewards were linked to people relying more on their internal signals (**Fig. 3A**), suggesting that a greater specialisation of neural responses to others’ rewards in this region occurs in people who are more aware and use interoceptive signals.

**Figure 3.**
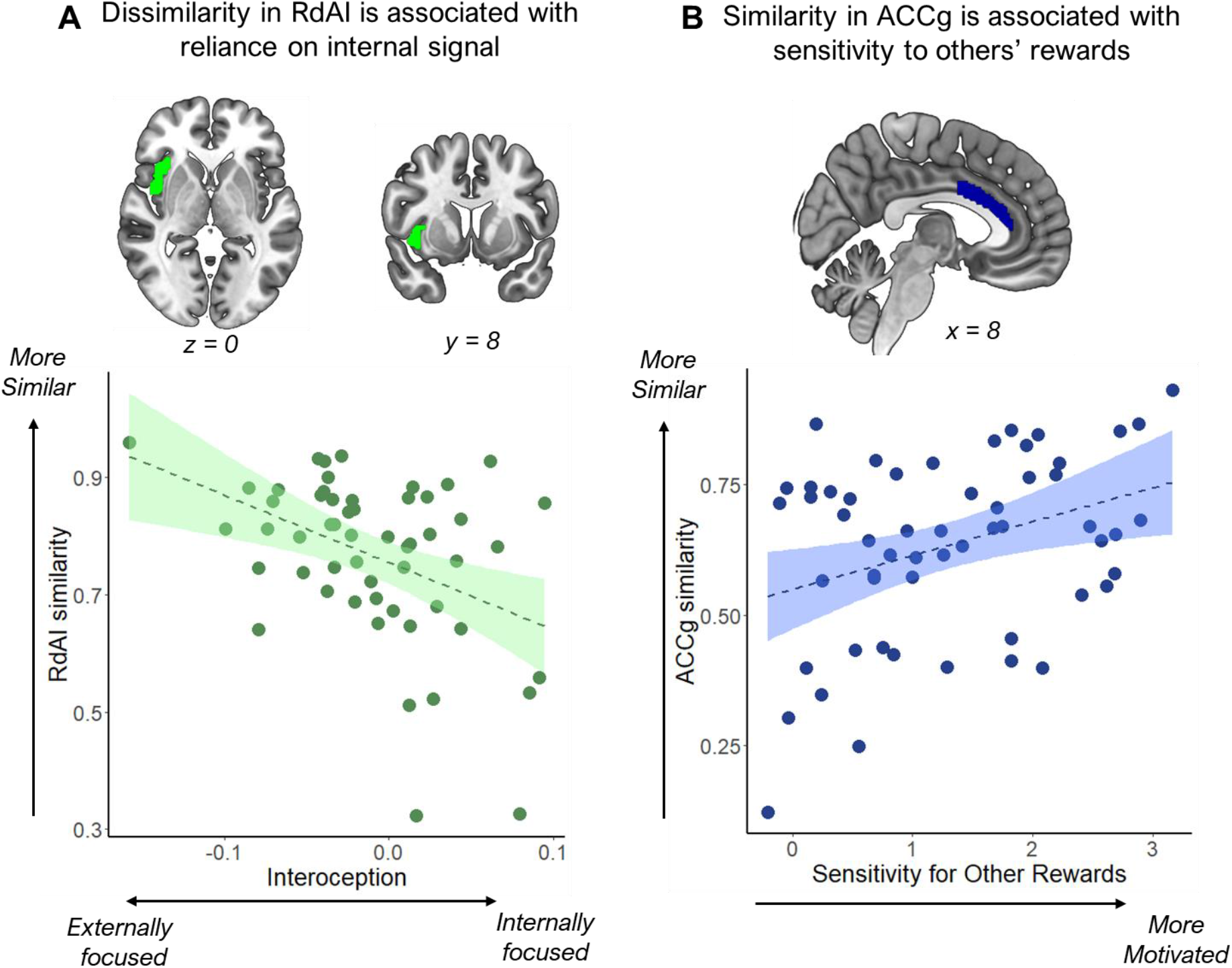
Similarity between self and vicarious reward responses in the right dorsal anterior insula (RdAI) and the gyrus portion of the anterior cingulate cortex (ACCg). A. Interoception was associated with multivariate similarity between self and other rewards only in the RdAI, among five ROIs (p < 0.008 FDR corrected). People who relied more on internal relative to external signals showed more dissimilar neural responses to reward between self and other. Y-axis depicts the similarity values, with higher values meaning more similarity between self and other reward representation. B. Prosocial motivation - specifically how incentivised a participant was to obtain rewards for others - was associated with multivariate similarity between self and other rewards only in the ACCg (p < 0.05 FDR corrected). Participants who were more motivated to work for others’ rewards showed more similar neural patterns between self and other rewards in the ACCg. X-axis corresponds to the reward betas obtained from a mixed model where decisions to work for others in the prosocial effort task was predicted by effort, reward and their interaction. Positive values indicate higher weights for others’ rewards.

Next, we hypothesised that variability in vicarious reward responses would be related to how incentivised a participant was by others’ rewards in the prosocial effort task. We used the same regression approach in each ROI, to test whether similarity in the neural response between self and other reward was related to a beta-weight reflecting someone’s sensitivity to others’ rewards in the prosocial effort task. We found a significant effect in only one ROI, the ACCg (b = 0.07, F_(1,53)_ = 7.36, p < 0.05 FDR corrected, **Fig. 3B**). Within the ACCg, greater similarity in responses between self and other reward was predictive of increased incentivisation by others’ rewards in the prosocial effort task. That is, people who showed a greater increase in choosing to help others as the rewards on offer increased, showed more similar neural patterns between self and other rewards. Note that both the results within the ACCg and RdAI were also present independently of the statistical method used to relate them, with similar results in both areas when Pearson correlation instead of regression models were used (see **Supplementary Information**). Furthermore, in both ACCg and RdAI, similarity between self and other was significantly above zero (**Supplementary Table S4**).

Analyses of the behavioural data revealed that interoception and sensitivity to others reward in the effort task were correlated, suggesting shared variance. However, analyses of the neural data demonstrated different effects of interoception and prosocial behaviour in distinct ROIs. In order to investigate whether the effects in the ACCg and RdAI remain when this shared variance is included in the model, two regression models were conducted, one in each region, in which the beta weight measuring incentivisation by other rewards from the effort task, and the interoception score, were both included as predictors of neural similarity. Both the effect of other reward in the ACCg (b = 0.06, F_(1,52)_ = 6.25, p < 0.02) and interoception in the RdAI remained significant (b = −1.21, F_(1,51)_ = 14.74, p < 0.001). As such, although people’s reliance on interoception and motivation by others’ rewards are related to each other, the responses in the ACCg and RdAI are linked specifically to prosocial motivation and interoception, respectively.

An advantage of using the prosocial effort task is that it also measures how sensitive people are to their own benefits (self reward), as well as how costly they find effort both when benefiting themselves (self effort) and others (other effort). To test the specificity of the effects above, we conducted two further regression models, one for the ACCg and one for the RdAI, including those three predictors. We found no significant effects in the ACCg nor in the RdAI for any of these predictors even at uncorrected thresholds (see **Supplementary Tables S4** and **S5** for details). Thus, vicarious rewards processing in the ACCg was specifically linked to how incentivised people were by others’ reward in the prosocial effort task, and similarity in the AI was linked specifically to interoception.

Taken together, vicarious reward signals in the RdAI and ACCg were linked to different processes: interoception, and motivation to obtain benefits for others, respectively. However, behaviourally, reliance on interoceptive signals was associated with higher motivation to obtain rewards for others in the effort task. Crucially, this link was specific for others’ rewards and not for self. Thus, the connectivity between RdAI (linked to interoception) and ACCg (linked to prosocial motivation) in reward processing might be stronger when evaluating others’ than self rewards. Could connectivity between ACCg and RdAI underlie this link between prosocial motivation and reliance on internal signals? We used beta series regressions (59) to assess individual differences in the functional connectivity between the ROIs in each task condition (self and other). Specifically, for each participant, we calculated the mean value for ACCg and RdAI during each trial and calculated the regression coefficient of those values across all trials and within each condition of interest. This allowed calculation of a beta estimate indicating the functional connectivity between ACCg and RdAI for each participant and condition, enabling comparison of the degree of connectivity between ACCg and RdAI in the self and the other condition. Significantly higher connectivity was found in the other compared with the self condition (t_(52)_ = 3.72, p < 0.001, **Supplementary Fig. S4**), suggesting that these areas are functionally more associated when reward events occurred to another person compared to participants themselves.

Next, we tested whether connectivity between RdAI and ACCg in response to other rewards was associated with reliance on interoceptive signals and/or sensitivity to others’ rewards in the prosocial effort task. Thus, a regression model in which the dependant variable was neural connectivity between these regions, having the interoception score and other reward betas in the prosocial effort task as predictors, did not reveal significant effects (see **Supplementary Table S6** for details).

## Discussion

We often observe good and bad outcomes for other people. However, how this experience of vicarious processing relates on our interoceptive signals or how willing we are to engage in prosocial behaviours remains largely unknown. Our results show that even when simply seeing rewarding outcomes for others, variability in vicarious processing in the ACCg and AI is linked to someone’s prosocial tendencies and their propensity to be aware and use interoceptive signals, respectively. In particular, whilst greater dissimilarity between neural responses to self and vicarious rewards was associated with increased reliance on interoceptive signals in the AI, greater similarity in the ACCg was linked to being more incentivised by rewards that could be obtained for others by prosocial acts. Functional connectivity between these regions was higher when participants observed others receiving reward outcomes than when they did themselves. Thus, these results indicate that processing of vicarious rewards is linked to levels of interoception and prosocial behaviour. In addition, at the behavioural level, how incentivised and motivated people are to obtain rewards for others relates to the propensity to be aware and use interoceptive cues, with more prosocial people being more focused on internal signals. This effect was specific, with reliance on internal signals not related to motivation to benefit oneself, or to the effort required to act prosocially. Taken together, these results suggest that variability in the patterns of neural responses to rewards are associated with interoception and prosocial behaviour.

The AI and ACCg have consistently been shown to be engaged when people passively monitor the receipt of rewards by either the self or others (2, 18–21), although this was not reflected in our univariate whole brain results. However, whilst both the AI and ACCg have been linked to vicarious reward processing, we show that variability in their response suggests a functional dissociation, which has the potential to unify separate lines of research into interoception, motivation and vicarious reward processing. Specifically, only the AI was associated with interoceptive signals. Previous research has suggested that the AI can be thought of as a secondary interoceptive cortex, due to the fact that it is heavily connected to regions that receive afferent signals from a number of sensory organs (34, 60–62). It has been argued that the function of AI is therefore to integrate interoceptive signals with other cognitive and affective processes (34, 60, 61). However, limited research has demonstrated a link between reward signals, that have often been found in the AI, to interoceptive processes. By using a multivariate technique, we were able to show that vicarious reward signals, specifically in the RdAI, are linked to reliance on interoceptive signals, highlighting that responses to vicarious rewards might reflect a stable trait that are associated to how much someone’s internal signals guide their behaviour.

Interestingly, the association between interoception and AI was specifically located in its dorsal part in the right hemisphere. This is consistent with previous work that proposes functional and anatomical division in the AI (63, 64). Thus, its ventral portion is believed to be mainly involved in affective reactivity toward salient outcomes impacting self and others, while its dorsal area (especially in the right hemisphere) has been more specifically linked to interoception, providing a bodily map for a wide range of mental processes (32, 34, 38, 46, 63–66). Posterior insula, involved in primary interoceptive representation of the physiological state of the body through thalamocortical pathways, has strong connections with dAI, and both share some of their connectivity fingerprints (34, 38, 46, 63, 65, 66). Through its connections with areas such as the prefrontal cortex, the ACC and the limbic system (34, 38, 46, 63, 64), dAI is crucial for self-awareness, where multiple sources of information are integrated to represent body-ownership and sense of agency (34, 38), which might be key for the distinction between self and other processes (67). Previous results indirectly suggest that the right AI, including its dorsal portion, have different mechanisms underlying vicarious and self outcomes (68–72), supporting the notion that RdAI might have representations of vicarious rewards anchored to bodily changes, important for self-other differentiation.

Notably, our results suggest that response to others’ reward in the ACCg better reflects people’s motivation to perform prosocial acts than their interoceptive propensity. Classical accounts have suggested that there is a functional dissociation between these regions, with the insula implicated in affective and bodily processes and frontal areas including the ACC implicated in motivating behaviour (34, 43–45). However, the presence of vicarious reward signals in both the AI and ACCg did not seem to fit with such a division. Here we show that it is the degree to which the AI represents others’ rewards as distinct from self rewards that is associated with interoceptive signals, and the same distinctiveness in ACCg that is associated with levels of motivation, specifically the willingness to help others. As such, the hypothesised division between AI and ACC suggested by classical accounts of their function is supported, with distinctiveness of vicarious signals of reward highlighting the distinct functions of the insula and ACC.

Although the classical account also posits that medial frontal cortex is involved in motivating behaviour, it is less clear that there would be a link specifically between vicarious signals in the ACCg and prosocial motivation. Previous studies in macaques have suggested that lesions that encompass the ACCg as well as surrounding portions of the ACC prevent monkeys from executing previously-learned prosocial behaviours, whilst leaving self-benefitting behaviours intact (73). Local field potentials and single-unit recordings have shown that neurons in the ACCg respond specifically when monkeys make a choice to deliver rewards to another rather than to themselves (74, 75). However, despite a number of studies showing vicarious reward signals in the ACCg, and that variability in these signals is linked to self-reported empathy and disrupted in autism spectrum disorders (20, 76, 77), very few studies in humans had linked these signals to directly-measured variability in prosocial behaviour. Here, we found that variability between people in the distinctiveness of ACCg vicarious reward signals was linked to how willing someone was to engage in prosocial acts, directly linking vicarious signals in this region with prosocial motivation.

Although these results in the ACCg and RdAI are correlational and require replication, they directly relate to broader discussions about whether common-currency or specialisation is more important for social and interoceptive processes (1, 25, 47, 67, 78). Usually, studies researching social information processing examine overlap between clusters responding to self or other rewards in a univariate analysis, or examine clusters that show a difference between self and other at the population level. Less work has examined individual differences in how distinct self and other reward processing is within a region, and there are very few studies that have used multivariate techniques that directly test how similar responses are between self and other rewards.

Recently, it has been demonstrated that multivariate pattern analysis techniques may be more robust for examining individual differences than traditional univariate statistical approaches (79). Here, by deploying this technique we showed that, within AI, specialisation for processing others’ rewards – less similarity in patterns between self and other reward – was related to greater reliance on interoceptive cues. As such, specialisation and identification of self and other rewards might be a distinguishing feature related to how AI processes interoceptive information. In contrast, in the ACCg, there was evidence that greater similarity in patterns between self and other reward was linked to increased prosociality in the effort task. Although this might argue in favour of a common-currency account, it is notable that the multivariate technique examines spatial disparity in terms of patterns, rather than overlap in voxels as examined with traditional methods. This work highlights that neither account in isolation can explain how the brain processes social information. This is also reflected in how the interaction between interoception and prosocial behaviour might be implemented in the brain. We showed that ACCg and RdAI are more functionally connected when people observe others’ reward outcomes than self. This could suggest that these areas, and potentially the link between interoception and prosocial behaviour, are mainly socially-specific. However, interpreting the links between behaviour, multivariate representations and connectivity seems too speculative, and conciliating these three approaches is beyond the scope of this study. Thus, the connectivity results add complexity to our main findings, and encourage future research to shed light on the ‘social brain’ debate. As such, common-currency and specialisation rooted accounts will need to be adapted to account for multivariate similarity analyses, and individual differences in such patterns, to move forward our understanding of the implementation of social information processing in the brain (47).

Finally, we found a behavioural link between prosocial motivation and interoception. Theoretical accounts posit that interoceptive signals might drive people to behave more prosocially (12, 16, 60, 80). Here, we show that people who have higher propensity to be aware and use interoceptive signals are more incentivised and motivated to help others, especially when the reward to be received by others increases, than people who rely more on external signals. Thus, we expand previous research reporting levels of altruism in economic games associated with levels of interoception (16). We used measurements that allow us to identify that increased sensitivity to rewards for others was specifically associated with reliance on interoceptive signal, rather than changes in one’s own experience of reward or effort. This might relate to the potential link between empathetic processes and interoception (81). Our results therefore support the notion that prosocial behaviour may be driven by how sensitive someone is to their internal bodily signals, mediated by sensitivity to others’ outcomes.

In conclusion, we show that vicarious reward processing, interoception and prosocial behaviour are closely linked. People who show a greater propensity to be aware and use interoceptive signals are more incentivised when they can obtain rewards for others and act more prosocially. Neural representations of passively observed vicarious reward in the AI and ACCg were able to predict people’s degree of reliance on internal signals and their levels of prosociality, respectively. These results suggest that social behaviour is complex and relies on both shared representations and self-other distinctions in the ACC and the AI, which might work together when facing a social situation. These findings highlight how important our everyday observations of others’ rewards may be for our subsequent internal and social lives.

## Materials and Methods

### Participants

The study was approved by the University of Oxford ethics committee. 80 healthy volunteers aged 18-38 years, recruited from the University and the city of Oxford, participated in the study, gave informed consent and were compensated for their time (£15 per hour for the fMRI session, £8 per hour for behavioural session, plus extra potential bonuses earned through the reward-related tasks). Participants were excluded if they had a history of neurological or neuropsychiatric disorders, psychoactive medication or drug use. Pregnant participants were also excluded from the study, as well as those who had participated in studies involving social interaction and who had studied psychology, addressing concerns that prior experience could influence psychological and brain processes.

61 participants (age M = 22.6, SD = 3.8, 32 females) were included in the behavioural analysis – four participants failed to show-up for the behavioural session, two participants expressed doubts about the social manipulation and the existence of the receiver, nine did not complete either the interoception respiratory task or the prosocial effort task, and four were excluded from analysis because of poor quality data in either of the two tasks (see below). From these 61 participants, 56 were included for the fMRI analysis (age M = 22.3, SD = 3.3, 30 females). One participant’s neuroimaging data was not registered due to technical issues, one participant showed excessive head motion, and three participants were excluded from analysis due to poor quality data in the value representation task (see below).

### General Procedure

Participants took part in an fMRI and a separate behavioural session. The value representation task was performed while in the fMRI scanner, whereas the interoception respiratory task and the prosocial effort task were tested as part of a behavioural testing session, performed at least one week after the fMRI session.

#### fMRI session: Value representation task

Before entering the scanner, participants were mock-randomly assigned the role of the Decider while another unknown person, a confederate, was assigned the role of the Receiver, following a well-established social manipulation protocol (8, 54) (see **Supplementary Information** for details). Participants were instructed that, on each trial, a computer would randomly select a value to add or to subtract to their total earnings (self trials) or to the earnings of the receiver (other trials). Thus, participants passively observed screens indicating that themselves or the anonymous receiver had gained or lost different amounts of reward. From this set of trials, participants were told that one of them would be randomly selected and added or subtracted to the receivers’ or to their own earnings at the end of the study. However, to avoid reducing participants’ compensation, a trial with a gain of £3 for themselves was always selected.

There were 12 different types of trials in the task according to a 2 (target: self, other) x 6 (value: −£15, −£9, −£3, £3, £9, £15) design. Each trial consisted of a screen indicating “You *OR* Receiver / Won *OR* Lost / Value” presented on a light grey background for 1400 ms followed by a 100 ms interstimulus interval. Furthermore, catch trials with the same duration were also presented to ensure attention to the task, as well as null trials presented for 3000 ms. The trial presentation order was pseudo-randomised using a “type 1 index 1” sequence which ensures that each stimulus is preceded and followed by all other stimuli (82, 83). For the purpose of this experiment, the four most efficient “type 1 index 1” sequences were chosen among 1000 randomly generated sequences using the Python code available online (https://cfn.upenn.edu/aguirre/wiki/public:t1i1_sequences/). Each of the four chosen sequences included 14 presentations of each stimulus – 14 repetitions x 14 trials (12 reward trials + one catch trial + one null/blank trial) - for a total of 196 trials. Since the full experiment included four sequences, 56 presentations of each stimulus were included with a scanning time of approximately 21 minutes across the four runs. The presentation order of the four sequences was counterbalanced across participants.

Catch trials were included to ensure that participants maintained attention across all trials without influencing the processing of the magnitude and the target of the gain or loss. During these catch trials, participants were asked to indicate as quickly as possible if recipient of the previous trial was the self or the receiver (i.e. “You or Receiver?”) or to identify the amount that was awarded in the previous trial compared to a randomly generated amount (e.g. “Won £15 or Lost £9”) using a response box. Accuracy at the catch trial task was computed by counting the number of correct answers and dividing it by the total number of catch trials. Participants who had an accuracy below 50% of the catch trials were excluded from analysis (n = 3).

#### Behavioural session: Prosocial effort task

The prosocial effort task measures how motivated participants are to obtain rewards for self and others (8, 48, 52). Participants in this task were instructed that, as Deciders, they would be paired with one of the Receivers of the experiment, but not the same one they had in the fMRI session. The task consisted of participants making decisions between options with different magnitudes of financial rewards (represented by number of credits) in exchange for different levels of physical effort (grip force). For each trial, participants chose between two options: a rest baseline option associated with no effort and low reward (1 credit); and a work offer option, which results in higher monetary gain (2, 4, 6, 8 and 10 credits) for higher effort that varies across 30, 40, 50, 60 or 70% of each participant’s maximum voluntary contraction (MVC). In half of the trials the reward was obtained by participants themselves (self trials), while in the other half rewards were received by the Receiver (other trials). If the work option was chosen, participants needed to make the effort required to obtain the credits on offer - they needed to squeeze a handle with the required force with their dominant hand for 1 s out of a 3 s window. They received zero credit if they failed to do so. If the rest option was chosen, participants rested for the same amount of time and obtained the single credit in exchange. The effort and reward levels in the work option varied independently over trials, and each effort–reward combination was repeated three times per beneficiary condition, giving a total of 150 trials: 75 self trials and 75 other trials. The rest option with one credit was used to make sure that there was a conscious and motivated decision to choose the alternative work option. If a choice was not selected, zero credits were given. All trials had the same duration, controlling for potential temporal discounting effects. Participants who did not actively choose any of the options for more than 10% of the trials were excluded from analysis (n = 1).

Before participants made decisions in the prosocial effort task, they were asked to grip a handheld dynamometer with as much force as they could to determine their MVC. Thus, the task measures similar effort levels across participants regardless of their variability in strength. After MVC estimation and prior to the main decision task, participants completed 18 trials where they experienced each effort level three times, and also learned to associate each level of effort with the elements in the pie chart (e.g. one element of the pie chart corresponded to 0% force, i.e. the rest option).

### Behavioural Session: Interoception respiratory task

The goal of this task (51) was to assess how much participants rely on their internal signals relative to exteroceptive cues when assessing the force of their exhalation. To assess participants’ force of exhalation, a standard peak flow meter was used. Participants who had a history of breathing difficulties did not undertake this task. On each trial, participants were first required to make a large exhalation into the peak flow meter. This first exhalation was taken as the ‘standard’ for that trial, i.e., 100% performance. Participants were then required to perform a second exhalation. In this second, ‘target’ exhalation, participants were told to aim to perform a percentage of the standard exhalation (e.g., 30%) for that trial. There were four possible targets for each trial - 30, 50 70 or 90% of the standard. Once the participant had performed the target, they were asked by the experimenter to estimate their performance as a percentage of the standard. The accuracy of their estimate (difference between their estimate of the force of their second blow as a percentage of the standard and the objectively-measured force of their second blow, see **equation 1**) served as the dependent variable for each condition.

Trials in the interoception respiratory task were completed under two conditions. In the *internal condition*, participants performed the standard and the target exhalations while hearing background white noise through headphones connected to a laptop (~79 decibels), ensuring they were unable to hear auditory cues produced by their exhalation, and thus could rely only on their internal signals when judging exhalation force. In the *external condition*, participants performed each exhalation with a background white noise coming from a laptop loudspeaker (~79 decibels) located approximately one meter from their right ear. Thus, in the external condition, although participants had an equally-distracting auditory input as in the internal condition, this auditory input did not mask the noise of their exhalation. As a result, participant could rely on auditory external cues when judging the force of their exhalation. For both conditions, the white noise started approximately one second before the exhalation, lasting for four seconds. Contrasting the performance in each of these conditions allows to measure the extent to which participants rely on interoceptive (relative to exteroceptive) signals. If in the external condition, where both internal and external signals are available to judge the force of one’s exhalation, a participant only uses external cues, then their performance is likely to differ markedly in the internal condition where external cues are unavailable. Conversely, if a participant relies entirely on internal signals to judge the force of their exhalation in the external condition – even though external cues are also available – then their performance should be relatively unaffected in the internal condition as internal signals are still available to be used.

During the task, and across conditions, participants were blindfolded, to prevent them using visual information to aid performance. For both internal and external conditions, participants completed six repetitions of the four targets presented in a random order, with a total of 24 trials per condition. The order in which the internal and the external conditions were presented was counterbalanced across participants. To ensure that the 30% target trial could be measured, participants were required to surpass a threshold of 200 L/min in their standard exhalation. If they did not accomplish this, the standard blow was repeated until the threshold was surpassed. No feedback about participants’ performance was provided across the experiment.

The peak-flow meter into which participants performed their exhalations was gently secured in a horizontal position using a vice clamp and elevated in line with each participant’s mouth using a stand. Participants were instructed to keep their hands resting at the bottom of the stand during exhalations, using their hands only to locate the gauge prior to performing exhalations, and to be still in the chair and sat upright, without pushing the mouthpiece forward while exhaling. Trials in which these conditions were not accomplished were repeated or removed from analysis. Participants with more than 10% of missed trials in either of the two conditions were excluded (n = 3).

### Analysis of behavioural data

#### Interoception score: reliance on internal vs external signals

To calculate participants’ reliance on their interoceptive signals, the absolute error scores were computed for each trial of the interoception respiratory task, such that:

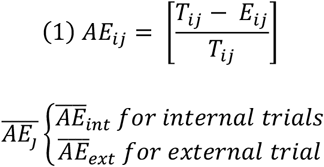

Where the absolute error *AE* in the trial *i* for the participant *j* is the absolute difference between the actual performance of the participant in target *T* as a percentage of the standard, and their estimation *E*, divided by their actual performance T. The mean of the AEs is computed separately for the internal and the external conditions for each participant. The interoception score was then the difference in performance between the internal and the external condition, such that:

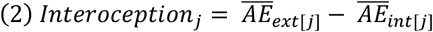

Where the interoception score for the participant j is the difference between their mean AEs in the external condition and in the internal condition. Scores below zero indicate that a participant made more mistakes in the internal than the external conditions, suggesting that they are more externally-focused. The interoception score therefore reflects how much participants rely on internal versus external signals.

#### Association between interoception and motivation

For the prosocial effort task, choices to work relative to rest were taken as an index of motivation to obtain self and other rewards. Mixed effects models were used to predict trial-by-trial decisions using the *glmer* function in R. Thus, two models were built to test whether interoception was linked to participants decisions, and to which variable (effort or reward for self or for other) in their motivation, such that:

Simple model

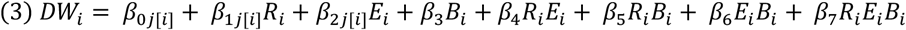

Interoception model

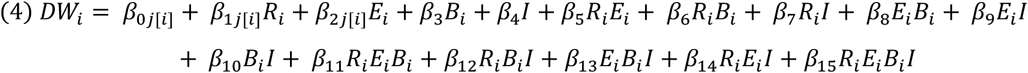

In the simple model, decision to work *DW* in the trial *i* was predicted by the fixed effects of reward *R*, effort *E*, beneficiary *B* and their interaction, with a random intercept clustered in each subject *j*. DW is a binary, factor variable. Random slopes for R and E were included as it is expected for these variables to vary across participants (8, 48, 52). The interoception model adds the interoception score *I* as a predictor together with its interaction with the other independent variables.

As a post-hoc analysis, we tested for Pearson correlations between the interoception scores and reward and effort beta values obtained for each participant from two mixed models where decisions to work were predicted by effort, rewards and their interaction. These reward and effort betas were used as a proxy for individual differences in participants’ sensitivities to reward and effort in self and prosocial decisions separately. Crucially, each model considered beneficiaries separately for self and other. These two models were clustered with random-intercepts for each participant, and had random-slopes for reward and effort, such that:

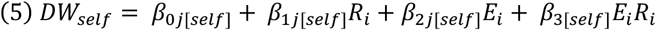

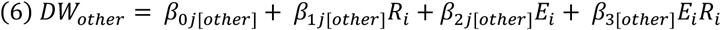

Where DW is specific for self and for other trials separately. Thus, the interoception score was correlated with motivation to work represented by participants’ beta estimates for reward and effort for self, other. For comparison between correlation coefficients Fisher Z-transformation were performed using the online tool at http://vassarstats.net/rdiff.html.

### fMRI acquisition and preprocessing

MRI data were acquired using a 3 Tesla Siemens Prisma MRI scanner. A structural scan was acquired at the start of the session using a magnetisation-prepared rapid gradient echo sequence with 192 slices; slice thickness: 1 mm; repetition time (TR): 1900 ms; echo time (TE): 97 ms; field of view (FOV): 192 × 192 mm; voxel size: 1 × 1 × 1 mm. Field map images were obtained immediately before the value representation task using a double-echo spoiled gradient echo sequence (TR: 488 ms; TE: 4.92/7.38 ms; voxel size: 3×3×3 mm; flip angle: 46°). Four runs of multiband T2*-weighted echo-planar imaging (EPI) volumes with BOLD contrast were collected during the value representation task (72 slices in interleaved ascending order; repetition time: 1570 ms; echo time: 30 ms; flip angle: 70°; field of view: 216×216 mm; matrix size: 108×108; voxel size: 2×2×2 mm with 1 mm gap; multi-band factor: 3; in-plane acceleration factor: 2). Each run was 5:46 minutes in length, during which 220 functional volumes were acquired.

Preprocessing was performed following a standard protocol using fMRIPrep 20.1.0rc1 (45), which is based on Nipype 1.4.2 (46). The details of preprocessing can be found in the Supplementary Material.

### fMRI analyses

#### First Level

First, a general linear model was created of the BOLD response at the participant level using SPM12, with regressors modelling the onset of each event, i.e. −£15, −£9, −£3, £3, £9, £15 for self and other, and the onset of the catch trials. The regressors were generated using delta functions convolved with SPM12’s canonical haemodynamic response function. This first level model also included nuisance regressors modelling non-neural sources: (i) 24 movement-related regressors comprising the six estimated head movement parameters (x, y, z, roll, pitch, yaw), their first temporal derivatives, their squares, and their squared derivatives; (ii) the first six components of the anatomical CompCor extracted using the CompCor method (84) implemented in *fMRIprep*; and (iii) a variable number of single event regressors indicating volumes exceeding a threshold of 0.5 mm framewise displacement or 1.5 standardised DVARS and considered as motion outliers (mean % of volumes that were motion outliers = 1.04%, SD = 0.91%). The four runs were concatenated in the model and constant regressors were added to model each of the four run’s mean. The null trials were used as the un-modelled baseline in all models. A high-pass filter with a cut-off of 128 s and SPM12’s *AR(1)* correction for serial correlation were applied during the model estimation.

To calculate functional connectivity between regions of interest in various conditions using beta series regressions (59), we also modelled the BOLD response for individual trials using the Least Square Single models approach (85). We therefore fitted a GLM for each trial of interest (468 trials) in which the first regressor was the onset of the trial of interest, the second regressor the onset of all other trials within the session and the other regressors were the nuisance regressors described above. To ensure that the single trial regressors included in the analyses were not collinear with nuisance regressors, we calculated the variance inflation factor (VIF) for each trial and excluded trials with a VIF > 4 from all analyses. This removed on average 3% (SD = 6%) of trials across participants.

#### Second level

For group statistical analysis, whole-brain SPM analysis was performed using a single-sample t-test to examine the contrast of self > other, and other > self conditions averaged across all reward events, using a cluster-level probability threshold of P_FWE_ < 0.05, with clusters defined by the voxel-level threshold P_uncorrected_ < 0.001.

#### Similarity analysis

We measured multivariate similarity between self and other trials by calculating the Pearson correlation between the average parametric maps for each condition of interest, i.e., self and other, in five ROIs. These correlation coefficients for each participant were used to assess the relationship between individual differences in self/other neural similarity and performance on the interoception respiratory task and the prosocial effort task. Null correlations between similarity in these ROIs and participants’ mean framewise displacement suggest that these values were not a product of participants’ movements in the scanner (see **Supplementary Information**).

We based our analyses on the AI and the ACCg as ROIs. For the AI portions, we used the parcellations created by Deen et al., 2011 (86) derived from a voxel-wise k-means clustering approach applied to resting-state. These parcellations divide the insular cortex in three different complexes for each hemisphere: posterior insula, dorsal AI and ventral AI. The 6 insula ROIs were acquired in 2 mm Montreal Neurological Institute (MNI) space directly from the author’s website (https://bendeen.com/data/). Within the AI there is putatively more than one anatomical zone, with a particular distinction in function between dorsal and ventral AI. However, it is unclear which of these sub-regions may be linked specifically to vicarious reward processing. As such, we used right and left vAI and dAI ROIs.

For the ACCg region, we used thresholded masks taken from the resting-state connectivity-based parcellation by Neubert et al., 2015 (87). This parcellation divides the frontal cortex into 21 different regions. The parcels were acquired in 2 mm MNI space directly from the author’s website (http://www.rbmars.dds.nl/CBPatlases.htm). We created the ACCg mask by modifying the original parcel using the *imcalc* function in SPM12. Thus, the left hemisphere parcel for the area 24ab was duplicated onto the right hemisphere to create a bilateral mask. Furthermore, those voxels located in the posterior portion of the mask were removed to capture more closely what the literature refers to as ACCg, corresponding to the gyral portion of the anterior mid-cingulate cortex (25, 88).

Similarity values obtained across these five ROIs were predicted by the interoception score and behavioural indexes in the prosocial effort task. For the latter, we used the reward and effort beta-weights obtained for each participant from the two mixed models previously described (**equations 5** and **6**) where decisions to work were predicted by effort, rewards and their interaction separately for self and other. For each regression model where ROI self/other similarity was predicted by behavioural measures, participants who had similarity values below and above three standard deviations were excluded. For the similarity values in RdAI, RvAI, LdAI and LvAI, robust regression models were used using the *rlm* function in R. For the similarity values in the ACCg, linear regression models were used instead as values in this region were normally distributed. All behavioural measures were normally distributed. Results obtained from these models were corrected for multiple comparison across the five ROIs using false discovery rate (FDR) (89, 90).

#### Functional connectivity analyses

We used beta series regressions (59) to assess individual differences in the functional connectivity between the ROIs in in each task condition. Specifically, for each participant, we calculated the mean value in ACCg and RdAI at each trial and calculated the regression coefficient between their combination across all trials for self and other. This allowed us to obtain a beta estimate parameter indicating the functional connectivity between ACCg and RdAI for each participant and condition. Paired t-test was used to test for differences in connectivity between self and other trials. Prior to this analysis, participants who had similarity values in both the ACCg and RdAI below and above three standard deviations were excluded (n = 3). Finally, a linear regression model was built having connectivity between RdAI and ACCg in other trials as a dependant variable, and the interoception score and other reward beta (**equation 6**) as predictors.

## Acknowledgements

This work was supported by a grant from the John Templeton Foundation (to MJC) and a Springboard Award (to MJC) from the Academy of Medical Sciences and the Wellcome Trust (SBF001\1008); a National Agency for Research and Development (ANID) DOCTORADO BECAS CHILE/2016 – 72170287 (to LSCH); and a Biotechnology and Biological Sciences Research Council David Phillips Fellowship (BB/R010668/1; BB/R010668/2) (to MAJA).

