## Supplementary material for "Neural representations of vicarious rewards are linked to interoception and prosocial behaviour": SI

### Supplementary Information for ‘Neural representations of vicarious rewards are linked to interoception and prosocial behaviour’

Luis Sebastian Contreras-Huerta, Michel-Pierre Coll, Geoffrey Bird, Hongbo Yu, Annayah Prosser, Patricia L. Lockwood, Jennifer Murphy, Molly J. Crockett, and Matthew A.J. Apps

**Supplementary Methods: Social manipulation procedure**

**Role assignment procedure in fMRI session.** Participants completed two kinds of trials in the fMRI and the prosocial effort tasks: self trials, where the beneficiary/recipient of the reward outcome was the participants themselves, and other trials, where the beneficiary/recipient of the outcome was another unknown person. We followed a rigorous protocol to manipulate the social aspect of the tasks performed by the participants to ensure they believed that the task outcomes affected real people. This protocol has been proven successful in making people believe and trust in the social consequences of other trials (1, 2).

This social manipulation protocol is framed as a *role assignment* procedure. Before participants went inside the scanner, they were told that two participants were taking part of the experiment, and that there were two roles involved which determine the characteristics of the tasks that each participant would perform. These roles corresponded to the Decider on one hand, and the Receiver on the other. The Decider would perform tasks in which they sometimes make decisions that could benefit themselves, and other times that could benefit the Receiver. The Receiver would perform a completely different set of tasks that do not benefit anyone. These roles are assigned randomly by selecting one of two balls from a box. However, and unbeknown to the participants, in reality all participants were assigned the role of the Decider, as the Receiver was a confederate. Once all the data were collected for each study, participants were contacted by email and debriefed about the deception used in the social manipulation.

The series of events to complete the protocol were the following. In the preparation stage, participants were inside the experimental room with the experimenter who explained the role assignment procedure. Then, the random assignment occurred, when a second experimenter arrived at the experimental room with the confederate standing on opposite sides of the participant in a semi-opened door, with the participant and confederate obscured from each other view. Both the participant and the confederate were instructed not to talk out loud, and they were handed a yellow rubber glove, preventing features that could reveal their identities from being seen. Once anonymity had been ensured, both participants and confederates were told to place their gloved hand in front of the door and to wave to one another in order to make sure that they were aware of the other person's presence. Then they proceeded to take a ball from a box held by one of the experimenters. The order in which they took the ball was decided by tossing a coin. Once the random assignment was completed, the second experimenter and the confederate left, and participants were informed that they would play the role of the Decider, according to the ball that they had chosen. Participants were informed that their choices in the tasks were anonymous, that they would not meet or interact with the Receiver, and that Receivers would perform a different unrelated task in parallel. These instructions aimed to minimise concerns about reputation, social desirability and reciprocity in their decision-making, all variables that can influence social behaviour (3, 4).

To ensure that participants believed in the social manipulation and the existence of the Receiver, participants were asked debriefing questions at the end of each session. These questions aimed to provoke participants to indicate their beliefs about the deception throughout the experiment. Crucially, these questions were subtle, and none of them directly ask the reliability of the procedure, avoiding inducing concerns about the veracity of the manipulation.

**Second social manipulation in behavioural session.** A second social manipulation was introduced in the behavioural session of the experiment. At the beginning of the prosocial effort task, participants received instructions and performed the calibration procedure (see **Materials and Methods** in the main text). Next, participants were introduced to the main decision task and its social aspects. Here, participants were told that they would be paired with one of the receivers of the previous fMRI session, but not necessarily the same one they had in their scanning session. A second experimenter then came to the room, and both the confederate/receiver and the participant stayed in opposite sides of a semi-open door, waiving their hands to each other, similar to the random assignment procedure in the fMRI session. After this, both the second experimenter and the receiver left, and participants proceeded to perform the prosocial effort task.

#### Supplementary Methods: fMRI preprocessing (this section was partly automatically generated by FMRIPrep)

**Anatomical data preprocessing.** The T1-weighted (T1w) image was corrected for intensity non-uniformity (INU) with N4BiasFieldCorrection (5), distributed with ANTs 2.2.0 (6), and used as T1w-reference throughout the workflow. The T1w-reference was then skull-stripped with a Nipype implementation of the antsBrainExtraction.sh workflow (from ANTs), using OASIS30ANTs as target template. Brain tissue segmentation of cerebrospinal fluid (CSF), white-matter (WM) and gray-matter (GM) was performed on the brain-extracted T1w using fast (FSL 5.0.9; 7). Volume-based spatial normalization to the MNI152NLin6Asym standard space was performed through nonlinear registration with antsRegistration (ANTs 2.2.0), using brain-extracted versions of both T1w reference and the T1w template. The following templates were selected for spatial normalisation: FSL's MNI ICBM 152 non-linear 6th Generation Asymmetric Average Brain Stereotaxic Registration Model (8).

**Functional data preprocessing.** For each of the four BOLD runs per subject, the following preprocessing was performed. First, a reference volume and its skull-stripped version were generated using a custom methodology of fMRIPrep. A deformation field to correct for susceptibility distortions was estimated based on a field map that was co-registered to the BOLD reference, using a custom workflow of fMRIPrep derived from Greve's epidewarp.fsl script and further improvements of HCP Pipelines (9). Based on the estimated susceptibility distortion, an unwarped BOLD reference was calculated for a more accurate co-registration with the anatomical reference. The BOLD reference was then co-registered to the T1w reference using bbregister (FreeSurfer) which implements boundary-based registration (10). Co-registration was configured with nine degrees of freedom to account for distortions remaining in the BOLD reference. Head-motion parameters with respect to the BOLD reference (transformation matrices, and six corresponding rotation and translation parameters) were estimated before any spatiotemporal filtering using mcflirt (FSL 5.0.9; 11). BOLD runs were slice-time corrected using 3dTshift from AFNI (12). The BOLD time-series were resampled onto their original, native space by applying a single, composite transform to correct for head-motion and susceptibility distortions. These resampled BOLD time-series will be referred to as preprocessed BOLD in original space, or just preprocessed BOLD. The BOLD time-series were resampled to MNI152NLin6Asym standard space and spatially smoothed using a 6mm full-width half maximum Gaussian kernel. Several confounding time-series were calculated based on the preprocessed BOLD: framewise displacement (FD), DVARS and three region-wise global signals. FD and DVARS are calculated for each functional run, both using their implementations in Nipype (following the definitions by Power et al., 2014, 13). The three global signals were extracted within the CSF, the WM, and the whole-brain masks. Additionally, a set of physiological regressors were extracted to allow for component-based noise correction (CompCor; 14). Principal components are estimated after high-pass filtering the preprocessed BOLD time-series (using a discrete cosine filter with 128s cut-off) for the two CompCor variants: temporal (tCompCor) and anatomical (aCompCor). Six tCompCor components are then calculated from the top 5% variable voxels within a mask covering the subcortical regions. This subcortical mask is obtained by heavily eroding the brain mask, which ensures it does not include cortical GM regions. For aCompCor, six components are calculated within the intersection of the aforementioned mask and the union of CSF and WM masks calculated in T1w space, after their projection to the native space of each functional run (using the inverse BOLD-to-T1w transformation).

##### Supplementary analyses for the association between behaviour and neural similarity

**Correlations between head movement parameters and neural similarity.** Spatial similarity between the representation of self and other rewards in ACCg, RdAI, RvAI, LvAI and LdAI were correlated with participants' framewise displacement motion parameter obtained from DVARS to test whether similarity values were influenced by participants' movement in the scanner. For each correlation between ROI self/other similarity and framewise displacement, participants who had similarity values below and above three standard deviations were excluded. Given that both the motion parameters and similarity values were not normally distributed, Spearman-rank tests were used for testing correlations. This analysis revealed that for all five ROIs, framewise displacement parameter was not significantly correlated with similarity values ($p > 0.05$).

**Correlations between behavioural indexes and neural similarity.** In addition to the regression models reported in the **Result** section of the main text, we also tested for correlations between the interoception score, motivational indexes (see **Materials and Methods, equations 5** and **6** in the main text), and similarity across the five ROIs. Pearson correlations were used to correlate brain similarity and behavioural measures. For each correlation between ROI similarity and behavioural measures, participants who had similarity values below and above three standard deviations were excluded. Furthermore, box-cox transformations were applied to those ROIs that showed deviation from normal distribution. All behavioural measures were normally distributed. Results obtained from these correlations were corrected for multiple comparison across the five ROIs using false discovery rate (FDR).

We first tested for correlations between the interoception score and similarity across the five ROIs. The only area that survived multiple comparison correction was the RdAI ($r = -0.41$, $p < 0.02$ FDR corrected), such that as people relied more on internal relative to external signals, they showed less similar neural patterns between self and other rewards. No effects were found in any of the other masks. Crucially, the effect of interoception on RdAI was significantly different from correlations between this area and all motivational indexes, i.e. sensitivity to effort and reward for self and other (Fisher-Z transformation, $p < 0.05$). Thus, the association between reward representations in the RdAI and interoception is specific, and different for self and other.

Next, we tested whether similarity in these five regions was associated with motivation. Specifically, beta weights for effort and reward separately for self and other were considered in this analysis. This analysis revealed that the only area that survived multiple comparison was ACCg, which was correlated with sensitivity to others' rewards ($r = 0.35$, $p < 0.05$ FDR corrected). Participants who showed more motivation to work for others depicted higher neural similarity between self and other rewards in this region. However, this effect was not significantly different from correlations between ACCg similarity and sensitivity to self rewards, nor with interoception. Yet, it was significantly different from correlations between ACCg similarity and effort betas (Fisher-Z transformation, $p < 0.05$). Taken together, these results are in line with those found using linear and robust regression models reported in the main text, suggesting an association between RdAI and interoception on one hand, and ACCg and prosocial motivation on the other.

#### Supplementary Figures

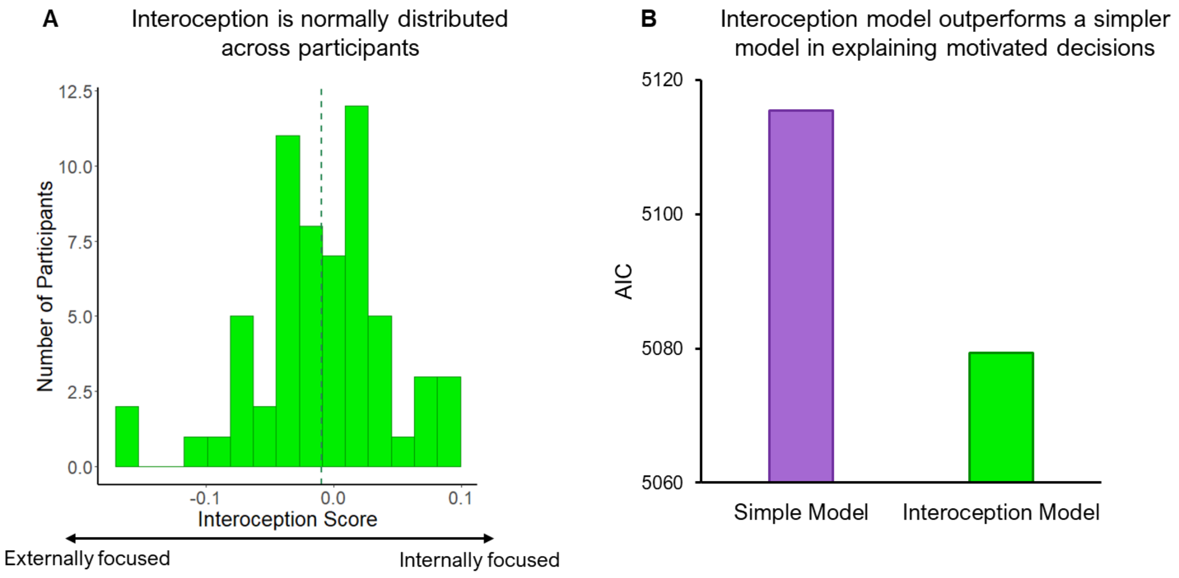


**Figure S1.** *A.* The interoception score followed a normal distribution with an average mean of -0.01 (indicated by the green dash-line). *B. Interoception score improved model fitting*. Adding interoception score as an independent variable predicting decisions to work in the prosocial effort task improved model fitting according to AICs (Akaike Information Criterion, y-axis). The simple model only included level of effort, reward and beneficiary as independent variables.

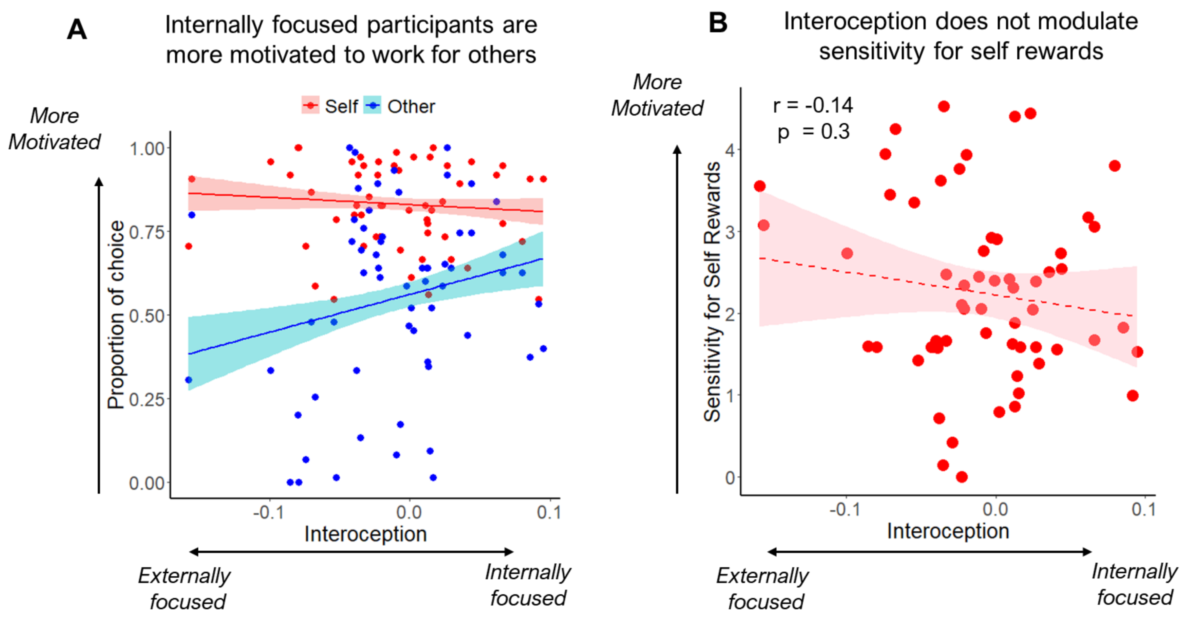


**Figure S2.** A. As participants relied more on internal than external cues in the interoception task (x-axis), they chose to work more to benefit others but not self in the prosocial effort task (y-axis). Shaded areas show the 70% confidence interval around the slopes. Individual points show the score of each participant for each condition. B. Null correlation between interoception (x-axis) and reward betas in the self condition (y-axis). Y-axis depicts the self betas from a mixed model predicting choices only for self. Shaded areas show the 95% confidence interval around the slopes. Individual points show the score of each participant.

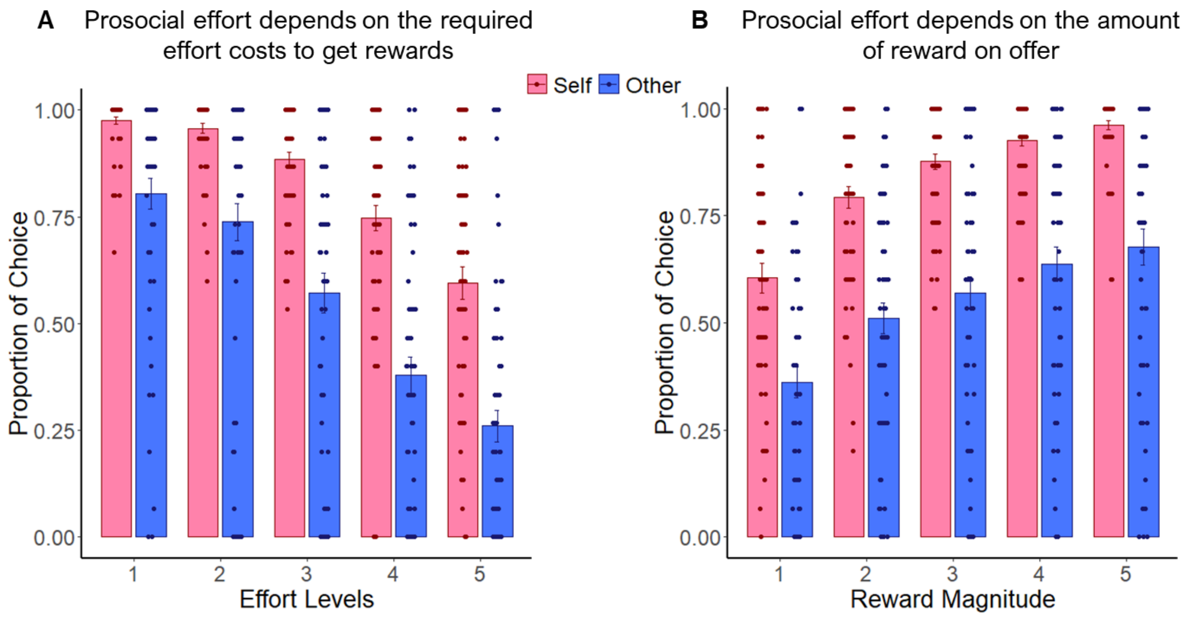


**Figure S3. Prosocial apathy effects are replicated.** *A.* In the prosocial effort task, participants were more reluctant to work at higher effort levels (x-axis), an effect that was augmented when the beneficiary was another unknown person relative to self. Y-axis depicts proportions of choosing the work offer (higher effort for higher reward) relative to a rest baseline option (no effort for small reward). *B.* Participants were more willing to work when the reward in offer was higher (x-axis), an effect that was amplified when the beneficiary was participants themselves compared with others. Y-axis depicts proportions of choosing the work offer relative to a rest baseline option. In all plots, dots depict each participant, and error bars represent SEM.

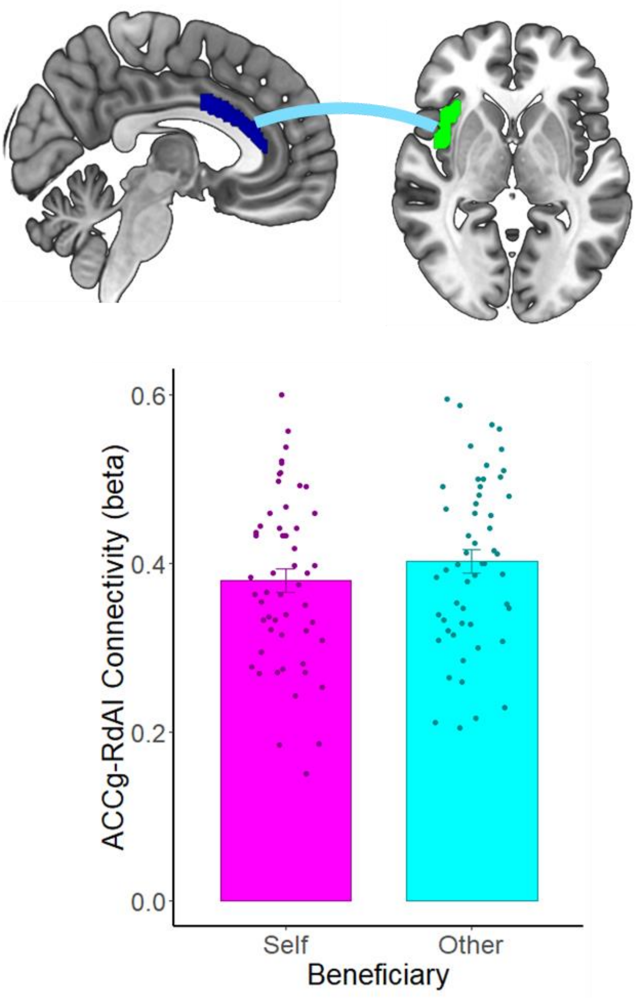


**Figure S4**. Connectivity between RdAI and ACCg is higher when processing reward events for others than for self. Dots depict each participant, and error bars represent SEM.

#### Supplementary Tables

**Table S1.** Results revealed by the interoception model, where trial-by-trial decisions to work or rest in the prosocial effort task were predicted by the effort level, the reward magnitude, the beneficiary of the reward, and participant's interoception score (for details see **Materials and Methods** section in the main text).

| *Effect* | *beta* | *SEM* | *z* | *p* |
|---|---|---|---|---|
| *Reward* | 2.44 | 0.21 | 11.81 | < 0.001 |
| *Beneficiary* | -3.59 | 0.13 | -27.47 | < 0.001 |
| *Effort* | -2.73 | 0.23 | -12.11 | < 0.001 |
| *Interoception* | -0.14 | 0.34 | -0.41 | 0.69 |
| *Reward*Beneficiary* | -1.04 | 0.12 | -9.04 | < 0.001 |
| *Reward*Effort* | -0.18 | 0.09 | -1.97 | 0.05 |
| *Beneficiary*Effort* | 0.39 | 0.11 | 3.37 | < 0.001 |
| *Reward*Interoception* | -0.1 | 0.2 | -0.48 | 0.63 |
| *Beneficiary*Interoception* | 0.65 | 0.12 | 5.3 | < 0.001 |
| *Effort*Interoception* | -0.21 | 0.22 | -0.96 | 0.34 |
| *Reward*Beneficiary*Effort* | 0.17 | 0.11 | 1.58 | 0.12 |
| *Reward*Beneficiary*Interoception* | 0.42 | 0.11 | 3.89 | < 0.001 |
| *Reward*Effort*Interoception* | 0.001 | 0.09 | 0.01 | 0.99 |
| *Beneficiary*Effort*Interoception* | -0.08 | 0.11 | -0.66 | 0.51 |
| *Reward*Beneficiary*Effort*Interoception* | 0.05 | 0.1 | 0.49 | 0.62 |

**Tables S2. and S3.** FWE-corrected results at the cluster level for univariate contrasts between self and other conditions.

*Table S2. Self > Other*

| *Regions* | *x* | *y* | *z* | *t* | *k* |
|---|---|---|---|---|---|
| *Lateral Occipital Cortex* | 40 | -80 | 6 | 5.03 | 307 |

*Table S3. Other > Self*

| *Effect* | *beta* | *SEM* | *z* | *t* | *k* |
|---|---|---|---|---|---|
| *Occipital Pole* | -6 | -88 | 0 | 14.3 | 16596 |
| *L Precentral Gyrus* | -54 | -6 | 48 | 5.04 | 427 |
| *Juxtapositional Lobule Cortex* | 4 | 0 | 64 | 6.36 | 394 |
| *R Precentral/Mid Frontal Gyrus* | 34 | -2 | 46 | 5.05 | 242 |
| *Superior Temporal Gyrus* | -56 | -10 | -6 | 4.96 | 155 |

**Tables S4 and S5.** Results revealed by two regression models where similarity between self and other rewards in the RdAI and the ACCg were predicted by sensitivity to self rewards, self effort and other effort in the prosocial effort task (betas extracted from **equations 5** and **6** described in the **Materials and Methods** section the main text)

*Table S4. Right dorsal anterior insula (RdAI)*

| *Effect* | *beta* | *SEM* | *F* | *p* |
|---|---|---|---|---|
| *Self Reward* | 0.02 | 0.02 | 1.74 | 0.19 |
| *Self Effort* | 0.01 | 0.02 | 0.07 | 0.79 |
| *Other Effort* | 0.01 | 0.02 | 0.07 | 0.8 |

*Table S5. Anterior cingulate cortex gyri*

| *Effect* | *beta* | *SEM* | *F* | *p* |
|---|---|---|---|---|
| *Self Reward* | 0.04 | 0.02 | 3.79 | 0.06 |
| *Self Effort* | -0.03 | 0.02 | 1.78 | 0.19 |
| *Other Effort* | -0.03 | 0.03 | 1.64 | 0.21 |

**Table S6**. Results revealed by a regression model were connectivity between RdAI and ACCg in the other reward condition was predicted by interoception and sensitivity to others' reward in the prosocial effort task (beta extracted from **equations 6** described in the **Materials and Methods** section in the main text)

| *Effect* | *beta* | *SEM* | *F* | *p* |
|---|---|---|---|---|
| *Other Reward* | -0.55 | 0.28 | 3.89 | 0.05 |
| *Interoception* | -0.001 | 0.01 | 0.01 | 0.92 |
